# Immune Cells Infiltration of Patient Derived Glioblastoma Cells spheroids in Acoustic Levitation in Bulk Acoustic Wave devices

**DOI:** 10.1101/2025.02.28.640539

**Authors:** Mousset Xavier, Kuermanbayi Shuake, Dupuis Chloé, Jeger-Madiot Nathan, Delaunay Virgile, Ibdaih Ahmed, El-Habr A. Elias, Chneiweiss Hervé, Junier Marie-Pierre, Aider Jean-Luc, Peyrin Jean-Michel

## Abstract

We describe an acoustofluidic device that allows scaffold-free structuration and culture of multi-cellular tumoroids composed of patient-derived glioblastoma cells only or in combination with non-cancerous cell. A PDMS chip of controlled height was created to allow acoustic levitation of cells using a 2 MHz transducer held on top of the chip. Cells are introduced into the chip through a dedicated inlet upstream of the resonant cavity. The specific design of the cavity together with the acoustic field allow the formation of tumoroids of cells in a precise and controlled manner within the levitation chamber. The acoustic and fluidic environment of the device was determined through experiments confronted with numerical simulations. The control of the flow within the chip was optimized to allow long-term culture of tumoroids and injection of cell culture media without disturbing the tumoroids in levitation. The tumoroids can be also structured, with sequential injections of the different cell types. Using microglia, we show that the acoustofluidic device allows the formation and culture in acoustic levitation of tumoroids mixing cancer cells with other cells populating the tumor as well as immune cell infiltration within the tumoroids. These results demonstrate the suitability of acoustofluidic levitation as an original 3D culture method adapted to the exploration of cancer growth at multiple levels.

## Introduction

Glioblastoma, the most common and aggressive malignant brain tumour, remains particularly resistant to current treatments, which typically combine surgical resection with chemotherapy and radiotherapy. Increasing evidence highlights the molecular and cellular heterogeneity of glioblastoma as key factors in treatment resistance. This heterogeneity results from the coexistence of distinct genomic clones, the exceptional plasticity of malignant cells, and the presence of normal cells co-opted by the tumour.

Components of the tumour microenvironment (TME) participate in promoting tumour growth and aggressiveness (Bikfalvi et al, 2023). The TME comprises a diverse set of neural cells (astrocytes, oligodendrocyte progenitors, neurons) and non-neuronal cells (immune cells, cellular components of blood vessels), as well as components of the extracellular matrix. Glioblastoma cells interact with the TME through various mechanisms, which influence both tumour progression and therapeutic resistance. For example, they induce tumour-associated macrophages (TAMs), the most abundant immune cellular component of glioblastoma, to adopt a pro-tumour state (Buonfiglioli, 2021). Similarly, glioblastoma cells interact with surrounding neuronal networks, creating a detrimental feedback loop that promotes tumour growth while disrupting neuronal activity (Venkataramani et al, 2022). A thorough understanding of these complex interactions between the tumour and the microenvironment is essential to progress towards more effective therapies. This requires the development of appropriate experimental models, simplified enough to allow a fine functional and molecular dynamic dissection of specific cellular interactions in a three-dimensional setting.

3D cell culture models of glioblastoma have generated substantial scientific interest due to their potential to better mimic the in vivo tumour environment. In contrast to conventional two-dimensional (2D) cultures, 3D models allow glioblastoma cells to grow in a more physiologically relevant context, preserving the complex architecture and cell-cell interactions found in actual tumours. These models facilitate the study of tumour heterogeneity, cell invasion, and resistance mechanisms in a controlled yet realistic setting. By incorporating elements like extracellular matrix components, 3D cultures provide insights into the influence of the tumour microenvironment on glioblastoma growth and therapy responses. They are particularly complementary of conventional 2D high-throughput drug screening, as they can better predict the efficacy and toxicity of potential treatments compared to 2D cultures. Overall, 3D cell culture models are a powerful tool for advancing our understanding of glioblastoma biology and for accelerating the development of more effective therapies.

The large majority of 3D models rely on matrix-based methods in order to constrain cells in a 3D setting with the help of a variety of hydrogels containing or not extra-cellular matrix components. Although relatively easy to establish, these models are limited by their reliance on scaffolds of uncertain impact on the cells. Organ-on-a-chip and cell bio-printing methods offer alternative 3D models including cell elements of the tumour microenvironment. They have notably allowed to expose tumour cell organoids to vascular-like structures (Wan, 2022, Tung, 2024). However, they are complex to implement and, for the bio-printing procedure, impose a strain on the cells. Most recent models have been developed based on ancient strategies grafting human glioblastoma cells in human nervous tissues obtained upon differentiation of human pluripotent stem cells (Nayernia, 2013) and now from human iPSC-derived brain organoids (Mangena, 2024). Finally, tissue from surgical resections of the patients’ tumours have been found to be amplifiable under constant rotation in culture (Jacob, 2020). These methods reproduce or conserve to a fair extent the tumour microenvironment but require months to be implemented or to access to fresh surgical material.

Acoustic levitation offers a novel and intriguing method of cell manipulation and cell culture, with several advantages over traditional techniques. By using sound waves to aggregate cells in a nutrient-rich medium (Bazou et al. 2012), acoustic levitation eliminates physical contact with a substrate, reducing stress and potential contamination. This method enables cells to grow in a three-dimensional environment with only cell-cell interactions, more closely mimicking the natural conditions within the body. Because cells are freed from the mechanical stress and cell-wall interactions inherent in 2D cultures, acoustic levitation can improve cell viability and function. This has been illustrated by the successful acoustic levitation culture of fragile cells such as mesenchymal stromal stem cells (Jeger-Madiot et al. 2021, Jeger-Madiot et al. 2022) or hepatic cells (Rabiet et al. 2024-1, Rabiet et al. 2024-2). Additionally, it allows precise control of cell positioning (Dron & Aider 2013, Jeger-Madiot et al. 2022), facilitating studies of cellular behaviour and interactions in a dynamic environment. This technique also holds promise for high-throughput screening (Sugier et al. 2023) and tissue engineering applications (Bouyer et al. 2016, Olofson et al. 2018, Tait et al 2019), as it enables the formation and culture of complex cellular structures and spheroids (Olofsson et al. 2021). Overall, acoustic levitation represents a cutting-edge and innovative approach to cell culture, with significant potential to advance research in cell biology and regenerative medicine (Rasouli et al. 2023).

In this study, we used acoustic levitation to rapidly and reproducibly grow tumoroids from patient-derived glioblastoma cells (GB-PDCs) and demonstrated the relevance of this approach to monitor the interaction of PDCs with immune elements in the microenvironment.

## Methods

### Acoustic Levitation

Acoustic levitation is used in this study to handle softly a large number of cells in well-defined positions inside the volume of a cavity where the cells ultimately can spontaneously self-organize into spheroids. It is also used to form more complex objects, using a second cell type that can be injected later. All these manipulations rely on the acoustic radiation force (ARF) applied to suspended objects in a resonant acoustic cavity (Bulk Acoustic Wave – BAW device). Indeed, if the resonance condition is satisfied (*h* = *n λ*_*ac*_/2, with *n* an integer and *λ*_*ac*_ the acoustic wavelength), a plane acoustic standing wave is generated inside the cavity (Fig. 1). In this case, every object in suspension in the fluid medium will undergo the axial component *F*^*rad*^ of the ARF. If the objects are considered as compressible spheres of diameter *d*_*p*_ small compared to the acoustic wavelength (*λ*_*ac*_ ≫ *d*_*p*_), then the ARF can be written as (Yosioka & Kawasima, 1955, Doinikov 1997, Bruus, 2012):

**Figure 1:**
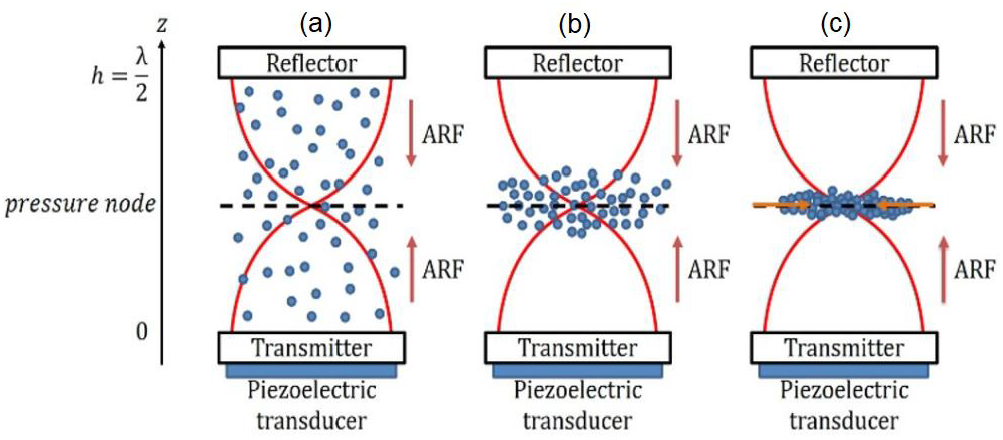
Schematic diagram of a *single-node* acoustic resonant cavity. Once the resonant condition is satisfied, an acoustic standing wave is generated inside the cavity (a), leading to the creation of an acoustic radiation force (ARF) whose axial component forces every suspending objects to move toward the acoustic pressure node (b). Once the objects have reached the pressure node, they remain trapped at this location, where they eventually form aggregates (because of the radial component of the ARF) and can be maintained in acoustic levitation as long as needed (c).

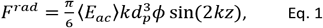

with ⟨*E*_*ac*_⟩ the time-averaged acoustic energy inside the cavity, *k* the acoustic wavenumber and *ϕ* the acoustic contrast factor, *z* being the axial axis. The acoustic contrast factor depends on the density and compressibility of the particles (*ρ*_*p*_, *β*_*p*_) and fluid (*ρ*_*f*_, *β*_*f*_) as:

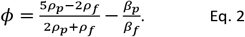

Objects with a positive contrast factor like cells lead to an axial ARF that will force them to move toward the acoustic pressure nodes. The acoustofluidics chips have been designed to respect the acoustic constraints, while searching for a geometry optimized to limit the flow perturbations and to allow gas transfers for cells incubations.

### Microfluidic Chip fabrication

The acoustofluidic chips were designed to control the flow over the tumoroids trapped in acoustic levitation in a cavity between the inlet and the outlet. Two types of acoustofluidic cavities were used. A chip was dedicated to microscopy imaging, meaning an observation of the lowest aggregate from under the chip using an inverted microscope. This chip is 40 mm long, 10 mm large and 1.5 mm high (containing up to 3 aggregates), with a total internal volume of 300 µL (Fig. 2A). The other chip was dedicated to cell culture in an incubator and side observations. The objective is to create and culture multiple aggregates for further biological analysis. They are also built to allow an observation from the side. This is useful to monitor the dynamic of self-organization of the aggregates over time. It is 40 mm long, 8 mm large and 5 mm high (containing up to 14 aggregates) with a total internal volume of 1mL (Fig. 2B).

**Figure 2:**
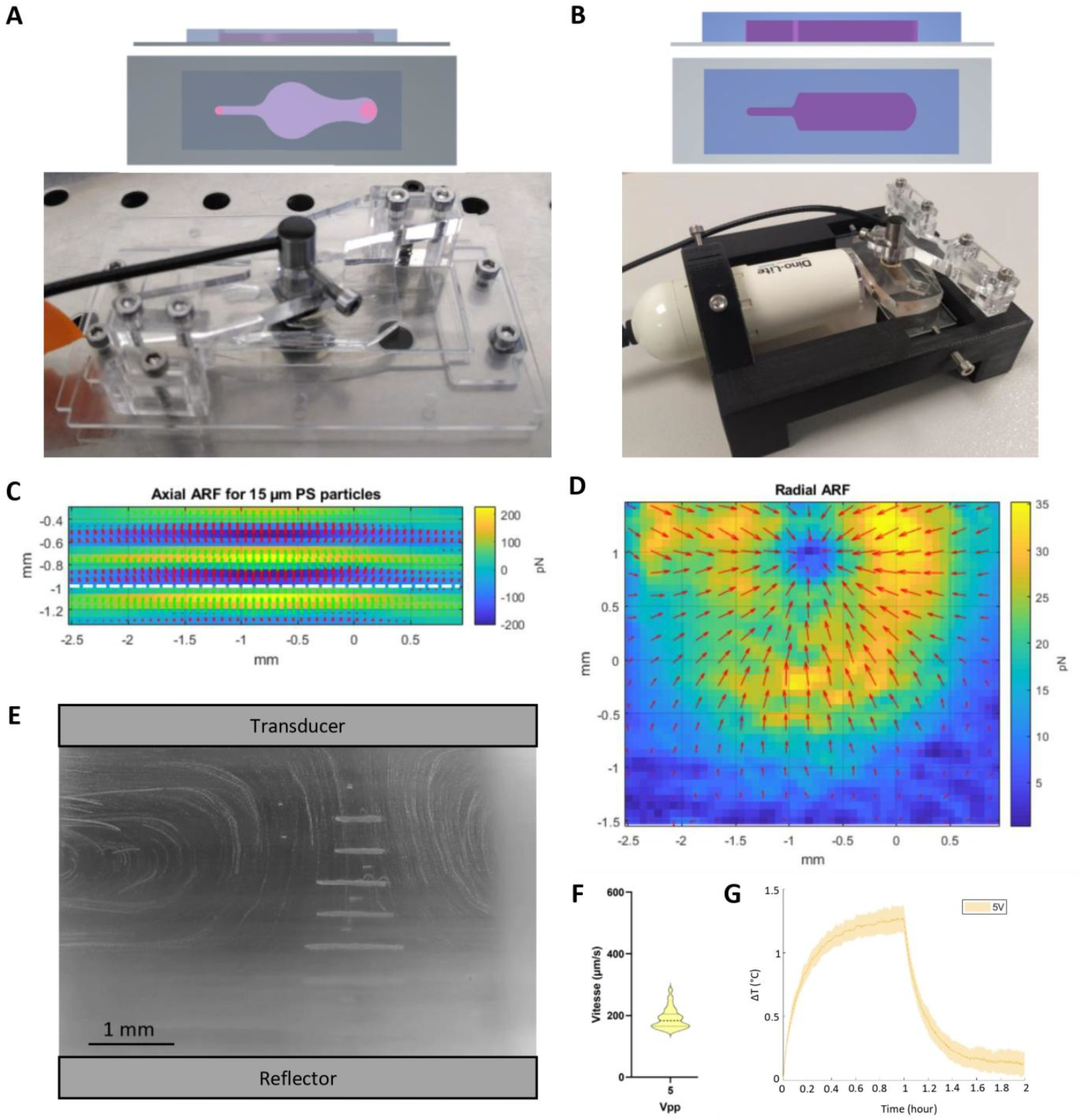
**A –** Schematic diagram of the *single-node* chip with the inlet and outlet openings (pink circles) asymmetrically designed for pipette handling. Below, picture of the *single-node* chip in culture configuration inside the PMMA holder is shown. The support and transducer holder were used to secure the chip and transducer together, facilitating handling of the system. The holder was designed to be placed as is on a microscope stage for direct imaging of levitated cells. **B –** Schematic diagram of the *multi-node* chip with below a picture of the complete system holder. The 3D printed support holds together the camera, the acoustofluidic chip and the transducer, thanks to integrated screws. This system enables easy and safe navigation between the incubator and the PSM without modifying the optical and/or acoustic settings, or the position of the chip. **C –** Computation of the ARF for the *multi-node* chip deduced from the acoustic pressure field according to the Gorkov’s theory (a<<λ) with a slice of the axial ARF component for a supplied tension of 5V. **D –** 2D field of the radial component of the ARF inside the levitation plane (white dashed line in Fig. 1C). **E –** Streaming visualization in a *multi-node* chip. The picture shown is a projection in time of the intensity maxima of a video at the beginning of a culture of GB-PDC R633 at 5V. The aggregates can be seen as thin sheet, indicating that the culture is still in its early stages. Streaming is similar for all cultures and does not depend on the particles, but on the acoustic parameters and chip dimensions. **F –** Speed of cells trapped in streaming measured on GB-PDC R633 at 2 MHz, 5V. Dotted line indicates mean and plain lines standard deviations. **G –** Variation of medium temperature as a function of time for different voltages applied with the piezoelectric transducer. Temperature measurements were carried out in a water-filled multi-node chip in a 37°C incubator. The thermocouple was placed in the levitation zone, below the transducer. Values are averaged from 2.00 to 2.03 MHz.

The acoustofluidic chips were fabricated using PDMS (poly-dimethylsiloxane), a gas-porous biocompatible material, using a protocol previously developed to grow neurons in microfluidic chips (Peyrin, 2011). Briefly, a master encoding the microfluidic chip was designed by laser cutting a 5 mm thick Plexiglas sheet and further glued on top of a polycarbonate petri dish (Greiner). The PDMS chip was made by pouring a degassed solution of 90% Sylgard™ 184 silicon elastomer base (Neyco) mixed with 10% curing agent (w/w) in the molding petri dish, with a thickness of at least 0.5 mm above the Plexiglas mold. The mold filled with PDMS was placed for 2 hours in an oven (Memmert) at 70°C. The PDMS imprint was cleaned with double sided tape (3M), and plasma bonded (plasma cleaner Diener electronic) on a 1 mm thick glass slide (VWR). The glass slide acts as a reflector for the ultrasound leading to stationary waves generation. Prior to plasma bonding, the glass slide was cleaned with precision wipes (Kimtech) soaked with isopropanol (VWR). After bonding, the chip was filled with water to preserve the hydrophilic properties of the PDMS acquired during plasma treatment. The chip and the PDMS caps were exposed 30 minutes to UV for sterilization, and stored at 4°C up to 15 days prior use.

### Acoustofluidic device

The acoustofluidic device consist of a PDMS/glass microchips and a 2MHz piezoelectric transducer (TR0205, Signal Processing™). The ultrasonic transducer is held by a home-made plexiglass holder for microscopy imaging of a single aggregate (Fig. 2A). A specific 3D-printed holder was designed to hold a digital USB microscope on the side of the *‘multi-node’* chip for the lateral imaging of the levitating tumoroids (Fig. 1B). The ultrasonic transducer was powered by an USB-powered digital arbitrary waveform generator (Handyscope HS5, Tiepie Engineering™). A thin layer of olive oil was used to ensure a good acoustic coupling between the transducer and the PDMS chip.

### Cell cultures

Patient-derived cells (PDCs), named 6240 and R633, were obtained from neurosurgical resection samples of distinct primary glioblastoma (El-Habr et al., 2017). The cells were expanded and cultured in conventional cell culture flasks in defined culture medium (human Neurocult NSA-H basal medium, StemCell) containing heparin (1 µg/mL, Sigma), EGF (20 ng/mL, AbCys SA), and bFGF (10 ng/mL, AbCys SA), as described (El-Habr et al., 2017). Human microglia clone 3 (HMC3) cell line (Russo 2018) was cultured in DMEM medium with 10% fetal bovine serum (Gibco). All cells were cultured in an incubator at 37 °C under 5% CO_2_. If necessary, HMC3 cells were labeled with the nontoxic Cell tracker red dye (Invitrogen). Cells were incubated for 5 min at 37 °C in PBS (500,000 cells/ml) containing the dye at 1 µg/ml before being washed with PBS and resuspended in fresh culture medium.

### Acoustofluidic spheroid formation, culture & retrieval

Acoustofluidic chips were mounted with the transducer under a Biosafety hood, and held in the Plexiglas holder. Olive oil was inserted between the transducer and the PDMS chip to allow for efficient acoustic coupling. Water inside the chip was replaced by cell culture medium. Single cell suspensions were prepared using Accutase (StemPro). A 50µL cell suspension was injected in the acoustofluidic device to reach a final concentration of 10^5^ cells/mL. It was then carefully mixed to homogenize the cells spatial distribution inside the levitation chamber.

The signal applied to the acoustic transducer was set at a frequency *f*_*ac*_ = 2 ± 0.05 MHz with a voltage amplitude of 7 V. These parameters maximize the efficiency of cell trapping in the levitation planes and their aggregation in the levitation planes. A good acoustic trapping also minimizes the risk of losing cells trapped in the pressure nodes during transport of the acoustofluidic chips from the hood to the incubator.

After injection of the cell culture medium with the cells and stabilization of the cells aggregates (initially layers of cells), the chip’s inlet and outlet holes were sealed with PDMS caps to prevent evaporation of the medium. The porosity of PDMS to gases allows exchanges between the medium and the air throughout the chip. Once the chips were installed inside the incubator, the voltage was decreased to 5 V for long-term culture. At the end of the experiment, the transducer is turned off and the tumoroids can be retrieved at the bottom of the chips.

### Characterisation of acoustic pressure field

Experimental measurements of the acoustic pressure amplitude inside the cavity were achieved using a Fiber Optic Hydrophone (Precision Acoustic™, UK) mounted on a XYZ translation stage. The sensor dimension is 10µm and its sensitivity is approximately 200mV/MPa. The chip was cut at the side to allow the introduction of the hydrophone inside the cavity, perpendicularly to the acoustic propagation axis (angular flat response of the sensor). The pressure field was scanned in a volume of 5 × 3 × 1 mm^3^ inside the cavity and with a spatial resolution of λ_ac_ /10 = 70 μm. The displacement of the probe was controlled by a dedicated software to probe automatically the volume of the cavity. The signal was recorded for 150µs with an acquisition frequency of 20MHz for each measurement points.

### Live cell imaging and Image analysis

The time-evolution of the size and shape of the cells aggregates created in the acoustic levitation planes was monitored using a USB digital microscope (AM2111, Dino-Lite™) with a resolution of 640 × 480 pixels, placed on the side of the chip. This small digital microscope was connected to a computer and controlled using the DinoCapture 2.0 software supplied by Dino-Lite. Snapshots of the cells were recorded every five-minutes throughout the length of cultures. A final image was recorded at the end of the experiment, just after turning off the transducer, once tumoroids had fallen at the bottom of the cavity. This image of the background without tumoroids was used in subsequent image processing. The images were processed using Matlab software (R2022a). Each image was converted to grayscale and binarized after background subtraction. Small objects were then removed and holes filled to improve detection of the parameters of interest. All detections were visually verified using a video file displaying the grayscale images along with the detected contour.

Three parameters were computed from the snapshots: the circularity (Cir), eccentricity (Ecc) and tumoroid diameter (D_eq_) defined respectively as:

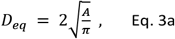

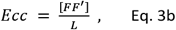

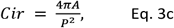

where (A) is the area and (P) the perimeter of the detected aggregate. [F F’] is the distance between each foci of the detected area considered as an ellipse and L is the length of its major axis. The diameter was normalized using its value at time t = 12 hours, when the cells layers had turned into spheroidal tumoroids.

The acoustic streaming was monitored with the measurement of the velocity field by Particle Image Velocimetry (PIV) using the PIVlab software (Thielicke, 2021; Jeger-Madiot et al, 2022) and cells as tracers.

### Tumoroid retrieval, cell counting and viability testing

After culture, levitated and non-levitated cells were retrieved from the chips in two different Eppendorf tubes and centrifuged at 3000 rpm for 1 min at room temperature. The tumoroids were exposed to Accutase (StemPro) for 5 min at 37°C and final cell dissociation was achieved with gentle up and down pipettings. Ten µL of cell suspension was mixed with an equal volume of Trypan Blue and incubated for 3 min at room temperature prior counting live and dead cells (Invitrogen Countess Automated Cell Counter, Thermofisher). The total number of counted cells ranged from 100 to 1000. Live/Dead staining was performed with LIVE/DEAD Cell Imaging Kit (ThermoFisher) according to the manufacturer instructions.

### Fixation, cryosections, tissue clearing and immunostaining

Tumoroids were fixed in 4% paraformaldehyde in PBS for 15 min at 4 °C, cryoprotected in 15% and 30% sucrose in PBS at 4 °C for 15 min and 30 min respectively, embedded in tissue freezing medium (Cryofix Gel, BIOGNOST), frozen in isopentane at – 40 °C and stored at – 80 °C until used. The tumoroids were cut into 25µm-thick cryostat slices. For immunostaining, the slices were washed with PBS for 15 min, and subsequently incubated in PBS containing 0.2% Triton (X100, Sigma) and 1% BSA (Sigma) for 5 min, and in PBS containing 10% BSA for 1 h at room temperature (RT). Primary antibodies were applied overnight at 4°C in PBS containing 5% BSA (anti-Ki67 1:1000, Agilent, M7240, A488-coupled anti-EGFR, 352907, BioLegend, 1:500), followed by rinsing and incubation for 2 hours at room temperature in PBS containing 5% BSA, 1:20000 DAPI (1050-A, Euromedex), and when relevant anti-mouse antibodies coupled to Alexa 633 (1/500, Invitrogen). After washing with PBS, slices were mounted using Mowiol (475904-M, Sigma) and stored at 4°C.

For tissue clearing, tumoroids were fixed in 4% paraformaldehyde in PBS for 15 min, followed by washing with PBS for 15 min. Tumoroids were then deposited into gene frame chambers (AB-0577, ThermoFisher) and adhered to superfrost slides (ThermoFisher). Slides were dried under the hood for 5 min and incubated for 2h at room temperature in permeabilization buffer containing 0.2% Triton and 20% DMSO. Tumoroids were then washed in PBS for 5 min, then once in 50% ethanol (VWR International) in PBS for 5 min, 80% ethanol in deionized water for 5 min, and finally in 100% dry ethanol for 5 min at RT. After removal of ethanol, tumoroids were covered with CytoVista Tissue Clearing Reagent (V11326, Invitrogen) 2h at RT. Then, CytoVista was removed and tumoroids were incubated with DAPI 1X overnight at RT, followed by washing twice with PBS 1X for 5 min each. Finally, tumoroids were covered with CytoVista and coverslips were added carefully on the gene frame chambers for imaging.

### Image Acquisition and Analysis

Immunofluorescence imaging was achieved using a Zeiss Axio-observer microscope and a Coolsnap HQ2 CCD camera, or a confocal microscope with Zen software (Zeiss LSM 980 Inverted, Confocal Laser Scanning Microscopy, Airyscan 2, Fluorescence-Lifetime Imaging module, laser diodes ranging from 405 nm to 639 nm, Colibri 7 LED light source, Zeiss ZEN Blue 3).

Acquired images were processed with FIJI (Schindelin, 2012), the StarDist plugin (Schmidt, 2018) and MATLAB. Cells were segmented relying on the DAPI staining using the StarDist plugin. The segmentation allowed to extract nucleus position and shape and to perform quantification.

Quantification of Ki67 was determined by averaging Ki67 fluorescent signal over the segmented nucleus or Region Of Interest (ROI). Positive or negative state of cell proliferation was determined by a threshold value defined visually for each image. Cell density was equal to the number of detected cells divided by the area of a circle with radius R0. The position of each nucleus positive for the staining was detected by thresholding the minima of the second derivative of the image. The position of the centroid of the tumoroid was calculated by averaging the different positions of the nuclei in the section, and the distance of each nucleus with regard to the centre was calculated (R/R0). The distribution of these positions was then plotted using a sliding box method with a box width of 30 µm.

### Statistical Methods

Statistical analyses were performed using unpaired t-test with Welch’s correction with Prism 6.0 software (GraphPad) and significance threshold set at p<0.05. Results are shown as means ± SD (Standard Deviation).

## Results

### Design and characterization of acoustofluidic cell culture devices

Acoustofluidic cell culture relies on the establishment of an ultrasound standing wave in a resonant cavity (Fig. 1) that must take into account fluidic control, biological constraints and geometries optimized to create an acoustic standing wave. In order to achieve long-term cultures of glioblastoma cells, we optimized two complementary configurations, one adapted to fluorescence microscopy (Fig. 2A) the other adapted to cell culture of many aggregates in an incubator (Fig. 2B). It is noteworthy that the microfluidic chip was designed to prevent media evaporation and bubble nucleation. Operating the transducer at 5V was found optimal to obtain efficient and reproducible trapping of GB-PDC in both single and multi-node chips. In the *multi-node chip*, the maximum amplitude of the measured acoustic pressure field was found to be about 300kPa, for a supplied voltage of 5V (see supplementary materials, movie 1). The acoustic velocity was deduced from the Euler equation in harmonic regime:

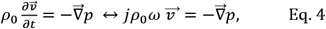

with 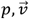 the acoustic pressure and velocity, *ρ*_0_ the fluid density, *ω* the angular frequency.

The acoustic energy density could also be evaluated using the relation 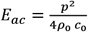 (Bruus 2012), leading to *E*_*ac*_ ≈ 10 *J*· *m*^−3^, a value comparable to other results found in previous studies on similar acoustic resonators (Bruus 2012, Dron & Aider 2013). To find a rough estimate of axial component of the ARF *F*_*rad*_ applied to the cells, the ARF acting on a polystyrene (PS) particle of a 15µm diameter has been calculated using the Gorkov’s theory (Bruus 2012). Indeed, the axial component of the ARF can be defined as the gradient of an acoustic potential:

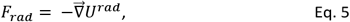

where *U*^*rad*^ is the acoustic potential defined as:

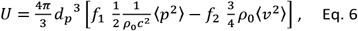

with 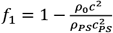 and 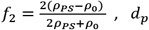, *d*_*p*_ the particle diameter, *ρ*_0_, *ρ*_*PS*_ respectively the densities of fluid and particle PS, *c, c*_*PS*_ the sound speed respectively in the fluid and polystyrene. Using these definitions, it is possible to derive the expression of the axial ARF given in Eq. 1 (Bruus, 2012).

Using these relations and knowing the properties of the fluid and polystyrene beads, the ARF field has been computed. The result obtained for a 15 μm polystyrene bead in a 1.5 mm high cavity is shown on Fig. 1C in a vertical plane. The axial ARF is maximum on the pressure antinodes with a value *F*_*rad*_ ≅ 200 *pN*. The particles or cells are then pushed by the axial ARF toward the pressure nodes (white dashed line in Fig. 2C). Once in the acoustic levitation plane (i.e. the pressure node) the axial component of the ARF vanishes, while the radial component *F*_*T*_ of the ARF is maximum. The 2D field of *F*_*T*_ in the horizontal levitation plane is shown in Fig 2D. One can see that the radial component of the ARF is oriented inward toward the cylindrical symmetry centre of the cavity, where the particles or cells will form an aggregate. The maximum value of the transversal component of the ARF is *F*_*T*_ ≅ 30 *pN*, much lower than the axial component, but strong enough to lead to the rapid formation of a large aggregate in acoustic levitation. In these conditions, it was possible to form large layers of PDC in acoustic levitation, that turned into tumoroids within 12h, with low risk of falling.

It should be noted that these calculations are valid only in these first steps of creation of large layers of cells. They cannot be applied to the large tumoroids that spontaneously form after 12 hours in acoustic levitation since the ≃ 300 µm final size of the tumoroid does not respect the Rayleigh approximation (d_p_ << λ_ac_). As they grow, tumoroids modify their interactions with the acoustic wave - and therefore the spatial organization of the acoustic field - but also their densities, both phenomena impacting the ARF. To ensure trapping of acoustically levitated tumoroids throughout the culture, ARF was applied at a minimum value of 5 V to provide a sufficiently powerful acoustic traps compensating for the weights of the tumoroids.

Acoustic wave propagation in liquid can trigger acoustic streaming (Mueller & Bruus 2014, 2015, Hoyos & Castro 2013, Castro & Hoyos 2016), which may be an issue to stabilize the aggregates over a long time. To quantify the streaming in the chips, the GB-PDC displacement in the vicinity of the levitation planes was monitored using the side observation of the multimode chip by the digital microscope. Time-series of snapshots were recorded, allowing the tracking of the cells (Video2) and an estimation of their velocities. The Fig 2E shows the trajectories of the cells injected into the chip. The large recirculations induced by the streaming can be clearly seen around stable aggregates in acoustic levitation in the center of the chip. Using cell tracking analyses it was possible to measure the average streaming velocity *v*_*st*_ = 200*μm* · *s*^−1^. The velocity distribution is shown in Fig 2F. Of note, the streaming does not perturb the levitation area where GB-PDC form very stable aggregates.

Finally, the temperature was measured using a thermocouple within the multi-node chip filled with cell media. The time evolution of temperature when the acoustics is turned on is shown in Fig. 2G. One can see a temperature increase of 1.25°C after one hour for these acoustic parameters. When the acoustics is turned off, the temperature decreases to reach its original value after one hour. To ensure a constant temperature of 37°C for cell culture, the temperature of the incubator was set at 36°C.

### Self-Organization of glioblastoma PDC under acoustic levitation

The self-organization of the layers of GB-PDCs into spheroids when cultivated in acoustic levitation was monitored at short and long times after cell injection in either the mono or the multimode chips. Cells were injected through the inlet port of the microchips while the acoustic transducer was activated (Fig. 2A, B). Aggregation at pressure nodes takes place sequentially after injecting the cell suspension into the chips. Injected cells are trapped within seconds as loosely packed planar layers (Fig. 3A) under the influence of the axial component of the ARF (Fig. 2C). This is followed by progressive packing of the cells over minutes at the center of the levitation plane (Fig 3A and B, Video1), or pressure node, under the action of the radial component of the ARF (Fig 2D) resulting in the formation of stable cell layers.

**Figure 3:**
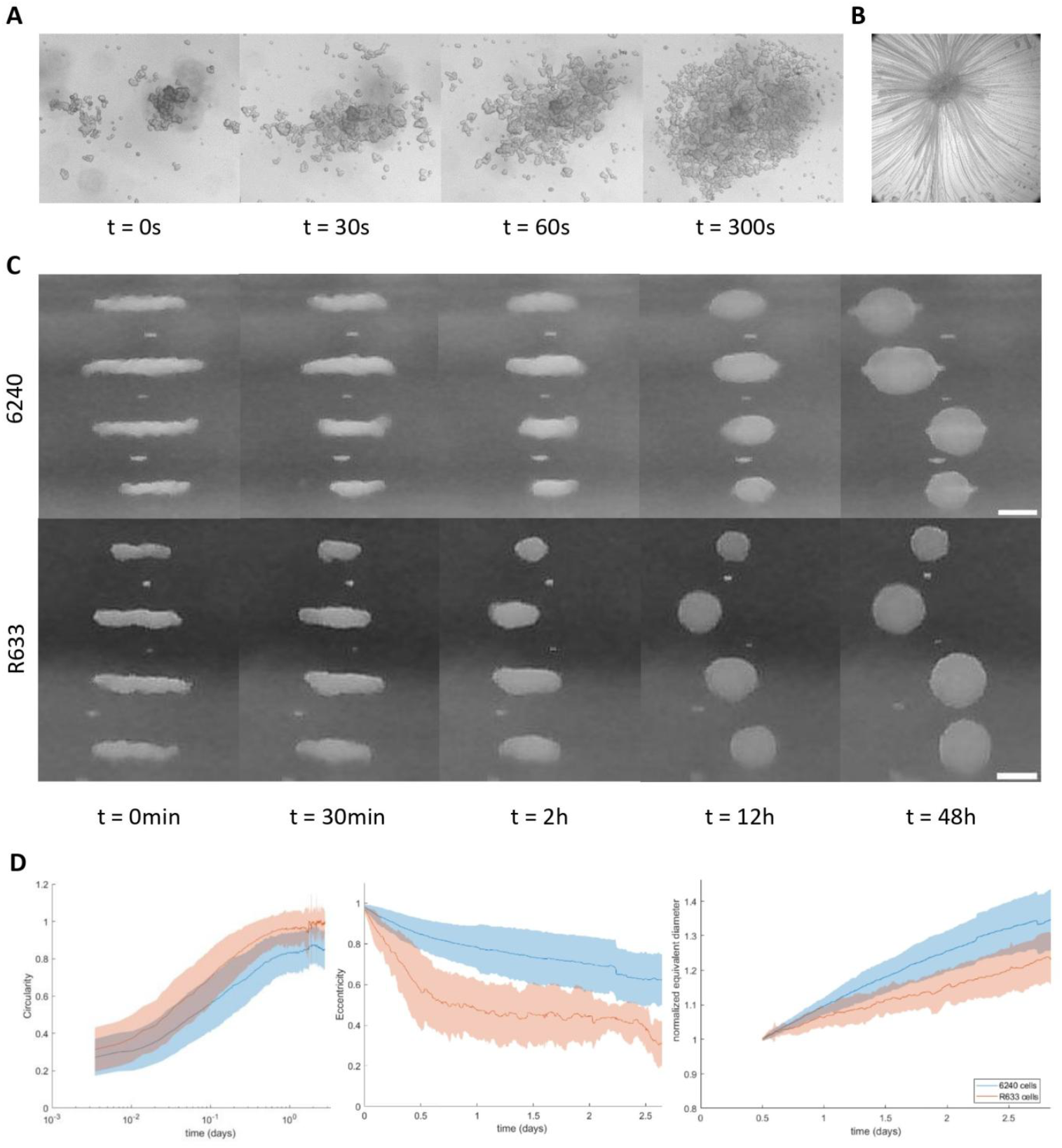
**A –** Aggregation of GB-PDC by acoustic levitation in a *single-node* chip. Cell aggregation takes place within minutes and is stable after 45 min. After this time point, almost no cell reach the aggregate. **B –** Time projection of intensity minima during cell cluster formation showing that GB-PDC aggregate from all directions. **C –** Side-view images of GB-PDC 6240 (top) and of GB-PDC R633 (bottom) tumoroids cultivated by acoustic levitation in a *multi-node* chip for 2 days. The various stages of spheroid formation up to 12h can be observed, followed by spheroid growth and size change between 12h and 2 days. Overall behaviour of the two cell lines was similar but differences in shape and speed of aggregation were observed with GB-PDC R633 forming tumoroids more rapidly and with a more spherical shape than GB-PDC 6240. Scale bar = 200 µm. **D –** Difference in shape according to cell line were observed as can be seen on circularity and eccentricity curves versus time for the two GB-PDC 6240 (n = 26) and R633 (n = 17), with measurements on different cultures and at different passages. As in C, there is a difference in spheroid formation that is reflected in the different eccentricity curves and distinct final circularity values. Evolution of tumoroid size by cell line is also shown by the equivalent diameter normalized by the equivalent diameter at 12 h for the same data set as circularity and eccentricity. As can be seen on the circularity and eccentricity plots, the shape of the tumoroid stabilizes after 12h in culture which is why the 12h time point was chosen for normalizing the tumoroid size.

We compared the behavior of two PDC lines (R633 and 6240) with different genomic backgrounds, different abilities to adhere to untreated culture vial surfaces, and distinct migratory properties (Saurty-Seerunghen 2022, Fig. S1A and B). PDC progressively self-organized over time into dense spheroidal aggregates with tumoroid-like shapes (Fig 3C). The kinetic of tumoroid formation was evaluated through the time evolution of the morphometric parameters, as defined in Methods section (see Equations 3a-c). The aggregates achieved roughly circular shapes (Cir = 1) after approximately 24 hours of culture (Fig 3D, left panel). Then the tumoroid size increased, suggesting active cells amplification inside the tumoroids (Fig. 1D, right panel). Both PDCs exhibit overall similar behavior in levitation, although cells with lesser migratory and adhesion properties tend to form more ellipsoidal tumoroids (Fig 3D, middle panel) while PDCs with highest adhesion properties self-organized more quickly into more spheroidal aggregates. Altogether these results show that PDCs self-organize in a contactless environment into tumoroid-like spheroids under acoustic levitation, regardless of their intrinsic specificities.

### Long term culture of PDC under acoustic levitation

Cell survival was evaluated after 6-7 days of culture by comparing cells captured in levitation to cells having developed under the form of small tumoroids outside of the acoustic field (Fig 4A). As shown in Fig 4B and C, cell survival assessed with trypan blue exclusion assay showed that levitation did not alter PDC survival as compared to control non-levitated cells, over 95% live cells being recovered in either condition. The cells were homogeneously distributed throughout the tumoroids with no evidence of a necrotic core, as shown with DNA labelling with DAPI and immunolabelling of EGFR, a transmembrane tyrosine kinase receptor overexpressed by glioblastoma cells (Fig 4D, left and middle panels). Cell densities were similar at both time-points evaluated (Fig 4E). Finally, we used Ki67 staining (Fig 4D, right panel) to determine whether PDC maintained active proliferation under acoustic levitation, as suggested by the increased size of the tumoroids over culture time. Quantification of Ki67-immunopositive cells showed that approximately 25% of PDC were undergoing proliferation at both time point evaluated (Fig 4F). Determination of the repartition of Ki67-immunoreactive cells within the tumoroids showed enrichment of proliferating cells in the outer layers of the tumoroids (Fig. 4G). Altogether these results demonstrate that acoustic levitation allows fast and efficient self-organization of glioblastoma PDC into actively proliferating tumoroids.

**Figure 4:**
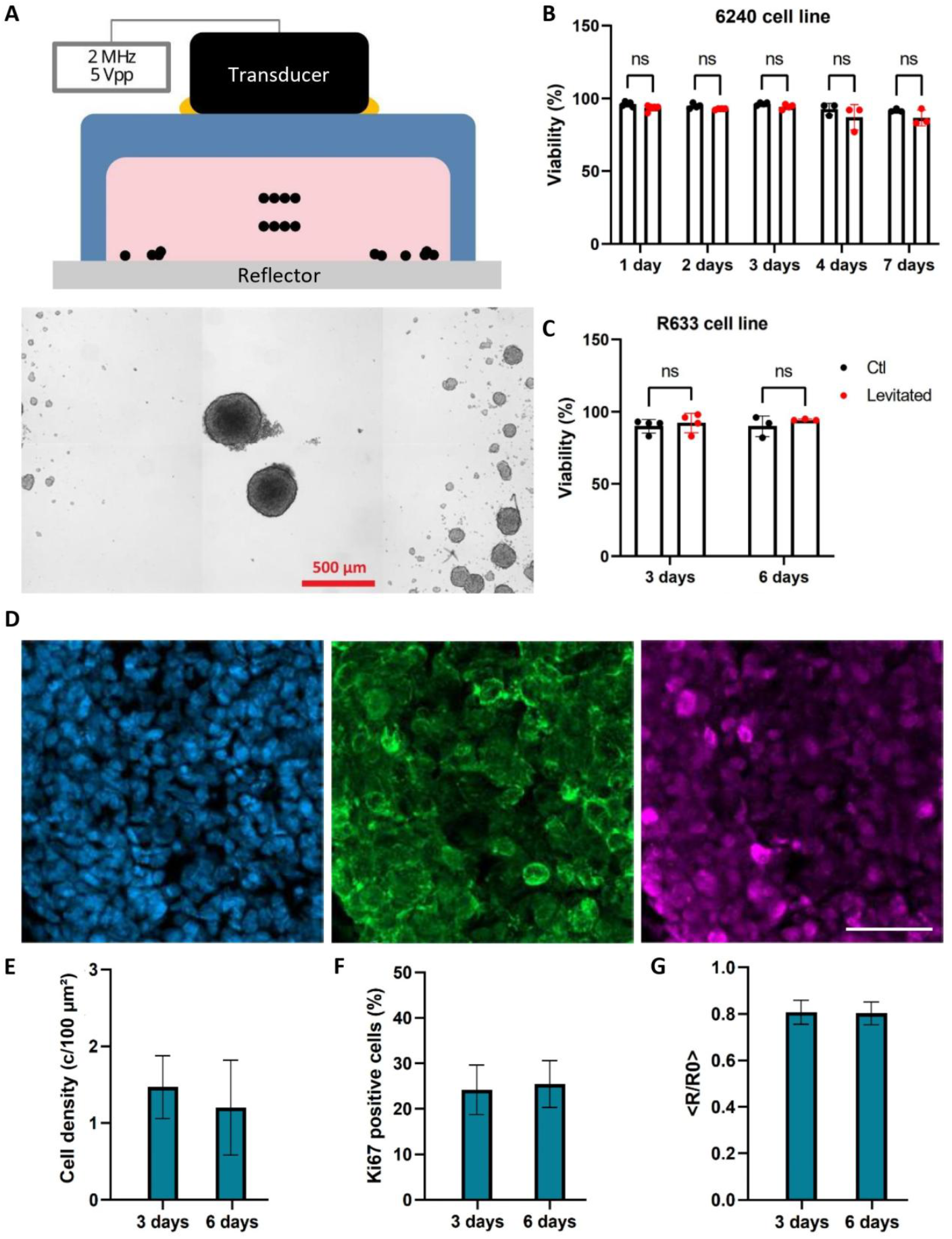
**A –** Schematic diagram of the *“multi-node”* chip (in blue) with the transducer (in black) containing the piezoelectric element on top of the chip, powered by a generator with a sinusoidal signal of frequency 2 MHz and power 5Vpp. The PDMS chip was bonded on top of a glass slide (in grey) which serves as reflector for the ultrasonic wave, producing the stationary field, and contains cell medium (in pink) and cells (black circles) in acoustic levitation, with some cells sedimented at the bottom of the chip. Different layers of cells aggregated at different pressure nodes are displayed. Below, picture showing a GB-PDC 6240 tumoroids culture after 5 days of culture at 5V, 2 MHz, with levitated tumoroids at the centre and control cells at the periphery having developed in contact with the glass slide outside of the acoustic field. **B and C**. Viability measurements by Trypan blue exclusion assay performed on PDC 6240 (B) and R633 (C) after 3 and 6 days of culture (mean±SD, n=3). Measurements were performed in different *single-node* chips at different passages. **D** Microphotographs of DAPI-labelled cell nuclei, EGFR-immunoreactive cells and Ki67-immunorractive cells, from left to right respectively. GB-PDC 6240 tumoroid cultured at 5 Vpp for 3 days. Scale bar = 50 µm. **E –** Cell density and **F –** Percentage of Ki67-positive 6240 GB-PDC at different time points in culture. **G –** Relative positions of Ki67-positive 6240 GB-PDC (R) from the centre of the tumoroid (R0) at different culture times.

### PDC and immune cell co-cultures in acoustic levitation

Next, we determined the suitability of acoustic levitation for implementing complex tumoroids that would integrate macrophage/microglia, i.e. non-malignant elements known to populate glioblastoma tumors, and to interact with malignant cells. The HMC3 microglia cell line was used, after verifying that its culture in PDC culture medium did not alter its survival and proliferation (Fig. S2). The tumoroids were first generated using a mixed cell suspension of PDC with human HMC3 microglia at a 10:1 ratio after staining microglia with a Cell Tracker Red membrane probe (Fig. 5A). Co-cultured GB-PDC/HMC3 tumoroids were cultured in multi-node acoustofluidic chips for 72 hours after injection of the mixed cell suspension. The patterning of the mixed cell suspension at acoustic pressure nodes was similar as the one of homogeneous PDC suspension. Fluorescent microscopy imaging of the resulting tumoroids showed patches of red labelled microglia at the periphery of and within the tumoroid (Fig. 5B). Tissue clearing and confocal imaging confirmed the presence of microglial cells inside the tumoroid masses (Fig. 5C), thus evidencing efficient generation of microglia/PDC mixed tumoroids.

**Figure 5:**
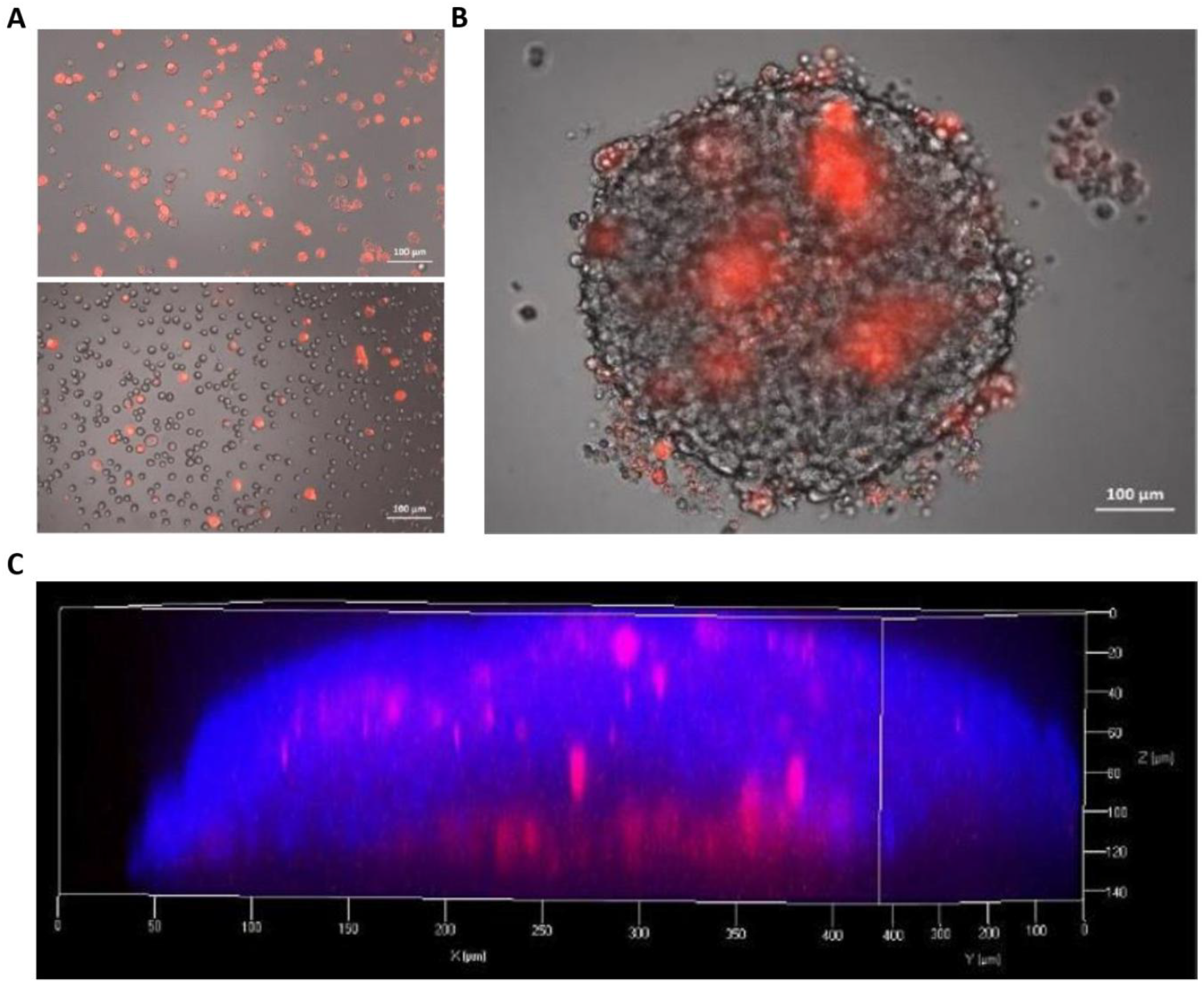
**A –** Fluorescent images showing HMC3 microglia stained with a Cell Tracker Red membrane probe (top panel) and a mixed cell suspension of PDC with human HMC3 microglia at a 10:1 ratio (bottom panel). **B –** Fluorescent image of the co-cultured GB-PDC/HMC3 tumoroid, with the two cell types mixed prior structuration, cultured in “multi-node” acoustofluidic chips for 72 hours demonstrating the efficient mixing of microglia and tumour cells with patches of red labelled microglia at the periphery of and within the tumoroid. **C –** Confocal image displaying the presence of microglial cells inside the tumoroid.

We next assessed whether acoustic levitation technique could be amenable to study tumor infiltration by microglial cells. We initially questioned the feasibility of structuring aggregates in concentric layers at the pressure nodes trough sequential injection of, first blue-, then white-colored polystyrene micro-beads (15µm-diameter, Fig. 6A). As expected, injection of the first bead population resulted in the formation of a stable and densely packed layer of microbeads at the pressure node as shown in Fig 6B1 (axial view from under the aggregate). Subsequent injection of the second bead population resulted in their progressive accumulation around the first layer of beads (Fig 6B2-3), through the action of the radial component of the ARF, ending in the formation of 2 concentric layers of beads (Fig 6B4). This protocol was implemented to inject red labelled microglial cells onto preformed PDC tumoroids maintained in acoustic levitation (Fig 6C). Lateral view imaging showed immediate trapping of microglial cells at pressure nodes trough the axial ARF component, prior to be progressively gathered around PDC tumoroids under the action of the transverse component of the ARF (Fig. 6D and video 3). The tumoroids were recovered after 2 more days of culture in levitation. Fluorescence microscopy imaging and confocal imaging after tissue clearing were used to verify the presence of microglial cells within the tumoroids. Microglia cells were found to have infiltrated the tumoroids, as shown in Fig. 6E and F.

**Figure 6:**
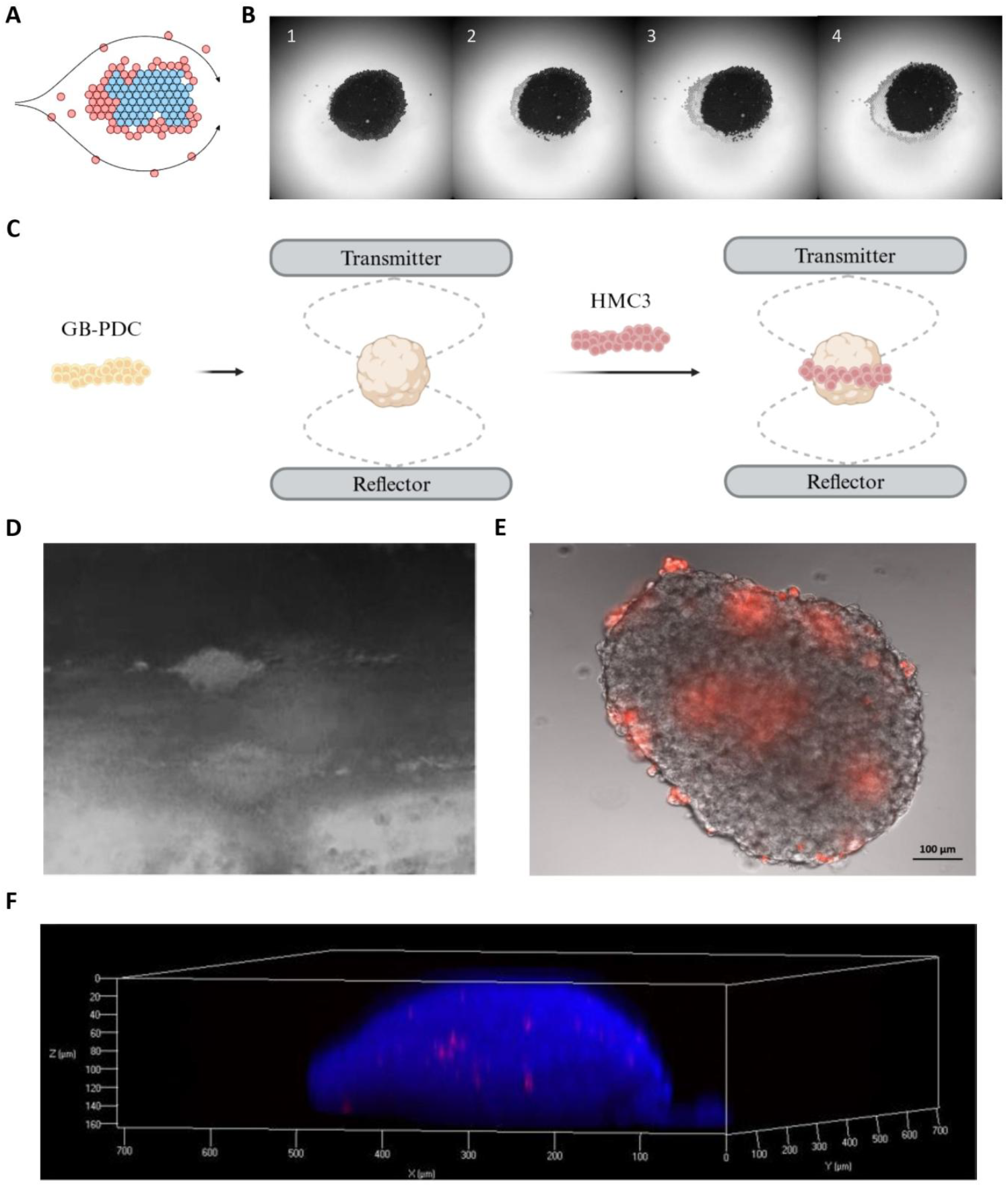
**A –** Diagram showing the combination of hydrodynamic and acoustofluidic effects allowing for the creation of a concentric structure with a first aggregate of blue particles being engulfed by red particles. Arrows correspond to the fluid trajectory. **B –** Images of blue polystyrene beads (in black) forming an aggregate (1) and further (2-4) surrounded by silica beads (in white). Both types of beads are 15 µm in diameter. **C –** Schematic of this phenomenon adapted for the infiltration of microglial HMC3 cells into previously GB-PDC tumoroids in acoustic levitation. Diagram created with BioRender. **D –** Picture of GB-PDC tumoroids trapped in acoustic levitation engulfed by HMC3 cells. **E –** Fluorescent image of the co-cultured GB-PDC/HMC3 tumoroid, with the GB-PDC structured in acoustic levitation 2 days prior infiltration by HMC3 cells, cultured in “multi-node” acoustofluidic chips for 72 hours demonstrating the efficient infilitration of microglia and tumour cells with patches of red labelled microglia at the periphery of and within the tumoroid. **F –** Confocal image displaying the presence of infiltrated microglial cells inside the tumoroid.

This ensemble of results demonstrates that acoustic levitation allows the formation of complex tumoroids gathering malignant and non-malignant cells, as found in patient tumors, thus offering a novel tool for exploring the dynamic interactions between malignant cells and their microenvironment.

## Discussion

Acoustic levitation represents an innovative approach to advance cell culture studies, particularly for challenging contexts like glioblastoma which need more realistic cell culture environments. In this study, we developed and characterized a microfluidic acoustic bioreactor capable of generating stable acoustic standing waves in a closed system, enabling precise control of the fluidic environment and facilitating long-term cell culture of many spheroids. The optimized microchip design allowed manual medium exchange without causing disruptive drag forces, streamlining the maintenance of cell cultures. Thus, we succeeded in performing long-term culture of 2 distinct patient-derived glioblastoma cell lines.

We demonstrated that malignant cells derived from the patients’ tumours cultured in acoustic levitation maintain active proliferation after self-organization into tumoroids that mirror the 3D morphology of tumour tissues. Notably, the system enabled the growth and three-dimensional self-organization of cells such as PDC-R633, which spontaneously grow under the form of 2D monolayers even in low-adhesion culture flasks. This underscores the potential of acoustic levitation to enforce rapid formation of tumoroids by promoting rapidly strong initial cell-cell contacts upon cell seeding. Interestingly, the spatial organization of the 2 distinct PDC lines presented subtle differences exemplified by self-organization of PDC-R633 into more ovoïd tumoroids. While there is no direct explanation, this may be linked to their distinct adhesion and/or migration properties as suggested by the observation that R633 cell lines strongly adhere and form migratory streams in conventional 2D cell cultures systems while 6240 cells spontaneously form cellular spheres in these conventional culture systems.

Furthermore, we leveraged this platform to model the intricate interactions between glioblastoma cells and other cell-intrinsic components of the tumour tissue. Microglia constitute an important fraction of the immune cell component of glioblastoma. While the role of microglia on glioblastoma cells is mostly modelled through 2D cocultures assays, 3D glioblastoma/microglia cell cocultures are technically challenging as both cells type have distinct adhesion properties (Niu et al 2023). By combining fluidic control with the transverse acoustic radiation force, our bioreactor facilitated the formation of controlled 3D co-cultures, enabling the study of microglia infiltration and interactions within a more physiologically relevant context. This advancement addresses a critical challenge in tumour immunology, where traditional 2D co-culture systems fail to replicate the distinct adhesion properties and spatial dynamics of immune cells and glioblastoma cells. Disposing of a simple model accessible to experimental studies of interactions between immune and glioblastoma cells is likely to offer valuable insights into tumor immunology and hence support the the development of immunotherapeutic approaches to combat this aggressive cancer. Easy retrieval of the tumoroids from the acoustic bioreactor enables indeed any type of analyses at either the tumoroid or the single cell level, including functional analyses on live cells. Structuration of tumoroids in acoustic levitation is in addition amenable to simulate varying microenvironmental pressures, for example by integrating additional immune cell types in various proportions, thereby enhancing the translational relevance of this model. Overall, our findings highlight the promise of acoustic levitation as a versatile tool for recreating complex tumour microenvironments. These insights open new avenues for exploring glioblastoma biology and notably the dynamics of interactions between malignant and normal cells that constitute this aggressive tumour.

## Conclusions

Acoustic levitation offers a novel and effective approach to advancing glioblastoma research by enabling the growth of patient-derived tumoroids and modelling complex immune-tumour interactions in a controlled, three-dimensional environment. The microfluidic acoustic bioreactor facilitates stable culture conditions, supports challenging cell lines, and provides a platform for studying microglia infiltration and intercellular dynamics with higher physiological relevance. This technology holds significant potential for improving our understanding of glioblastoma biology and accelerating the development of therapeutic strategies.

## Author contributions

Conceptualization, JMP, JLA, XM, EEH. Methodology, XM, JLA, JMP, EEH, MPJ, HC. Experimental investigations, XM, SK, CD, NJM. Data analyses, XM, SK, CD, NJM, EEH, MPJ, JLA, JMP; Writing – Original Draft, JMP, XM, MPJ – Review & Editing, all. Project Administration, JLA, HC. Funding Acquisition, JLA, HC, MPJ.

## Data availability

Data will be made available upon request

## Acknowledgement

We thank the BRIGHT research team from the ICM institute for providing us with the HMC3 cell line.

